# Reassessing associations between white matter and behaviour with multimodal microstructural imaging

**DOI:** 10.1101/2020.12.15.422826

**Authors:** Alberto Lazari, Piergiorgio Salvan, Michiel Cottaar, Daniel Papp, Olof Jens van der Werf, Ainslie Johnstone, Zeena-Britt Sanders, Cassandra Sampaio-Baptista, Nicole Eichert, Kentaro Miyamoto, Anderson Winkler, Martina F. Callaghan, Thomas E. Nichols, Charlotte J Stagg, Matthew Rushworth, Lennart Verhagen, Heidi Johansen-Berg

**Affiliations:** Wellcome Centre for Integrative Neuroimaging, FMRIB, Nuffield Department of Clinical Neurosciences, University of Oxford; Oxford Centre for Human Brain Activity, Wellcome Centre for Integrative Neuroimaging, Department of Psychiatry, University of Oxford; Section Brain Stimulation and Cognition, Department of Cognitive Neuroscience, Faculty of Psychology and Neuroscience, Maastricht University, Maastricht, The Netherlands; Maastricht Brain Imaging Centre (MBIC), Maastricht University, Maastricht, The Netherlands; Department of Clinical and Movement Neuroscience, Institute of Neurology, University College London, WC1N 3BG; Wellcome Centre for Integrative Neuroimaging, Department of Experimental Psychology, University of Oxford; National Institute of Mental Health, National of Health, Bethesda, MD, USA; Wellcome Centre for Human Neuroimaging, UCL Queen Square Institute of Neurology, UCL, London, UK; Oxford Big Data Institute, Li Ka Shing Centre for Health Information and Discovery, Nuffield Department of Population Health, University of Oxford; MRC Brain Network Dynamics Unit, University of Oxford, Oxford, OX1 3TH, UK; Donders Institute for Brain, Cognition and Behaviour, Radboud University Nijmegen, Nijmegen, Netherlands

## Abstract

Several studies have established specific relationships between White Matter (WM) and behaviour. However, these studies have typically focussed on fractional anisotropy (FA), a neuroimaging metric that is sensitive to multiple tissue properties, making it difficult to identify what biological aspects of WM may drive such relationships. Here, we carry out a pre-registered assessment of WM-behaviour relationships in 50 healthy individuals across multiple behavioural and anatomical domains, and complementing FA with myelin-sensitive quantitative MR modalities (MT, R1, R2*).

Surprisingly, we only find support for predicted relationships between FA and behaviour in one of three pre-registered tests. For one behavioural domain, where we failed to detect an FA-behaviour correlation, we instead find evidence for a correlation between behaviour and R1. This hints that multimodal approaches are able to identify a wider range of WM-behaviour relationships than focusing on FA alone.

To test whether a common biological substrate such as myelin underlies WM-behaviour relationships, we then ran joint multimodal analyses, combining across all MRI parameters considered. No significant multimodal signatures were found and power analyses suggested that sample sizes of 40 to 200 may be required to detect such joint multimodal effects, depending on the task being considered.

These results demonstrate that FA-behaviour relationships from the literature can be replicated, but may not be easily generalisable across domains. Instead, multimodal microstructural imaging may be best placed to detect a wider range of WM-behaviour relationships, as different MRI modalities provide distinct biological sensitivities. Our findings highlight a broad heterogeneity in WM’s relationship with behaviour, suggesting that variable biological effects may be shaping their interaction.

**Highlights:**

- Pre-registered testing of microstructural imaging across modalities (FA, MT, R1, R2*) to test WM-behaviour relationships.
- Partial support for FA-behaviour relationships hypothesised based on previous literature.
- Multimodal approaches can help detect WM-behaviour relationships that are not detected with FA alone.
- Sample sizes of 40 to 200 may be needed to detect myelin-behaviour relationships in joint multimodal analyses.
- Variable biological effects may be shaping WM-behaviour relationships.

## Introduction

The past decade has shown that White Matter (WM), and in particular the myelinated structures that dominate it, have more varied functions than previously thought, from trophic support of axons (Fünfschilling et al., 2012; Nave, 2010) to active regulation of physiological and behavioural processes (Kaller et al., 2017; Lazari et al., 2018; Steadman et al., 2019). These basic biology findings suggest that WM may play a role in brain physiology and behaviour, and that WM could be targetted for therapeutic gain in neuropsychiatric disorders (Gibson et al., 2018; Vanes et al., 2020).

In humans, much evidence on the role of WM has come from a large body of studies linking behaviour to diffusion-tensor-based metrics such as fractional anisotropy (FA), a metric derived from diffusion weighted imaging that is sensitive to features of WM microstructure (Boekel et al., 2015; Johansen-Berg, 2010; Lazari and Lipp, 2020; Roberts et al., 2013). While these studies have provided seminal evidence for a link between WM and human behaviour, questions remain about the generalizability and interpretation of these effects.

FA-behaviour relationships are particularly difficult to interpret on a biological level. Diffusion signals are sensitive to a broad range of tissue properties, including myelination levels, fiber orientation, axon diameter, astrocyte and vascular morphology (Farquharson et al., 2013; Sampaio-Baptista and Johansen-Berg, 2017; Stolp et al., 2018). Therefore, a given FA-behaviour correlation could arise from a diversity of microstructural patterns (Zatorre et al., 2012). Moreover, while other tensor-based metrics can be derived from diffusion-weighted imaging, it is unclear whether they differ from FA in their biological sensitivity (Lazari and Lipp, 2020).

In recent years, an increasing number of techniques (Figure 1) have been successfully applied to the study of WM, and of WM myelination in particular (Heath et al., 2018). As WM is dominated by myelinating oligodendrocytes, many of these techniques have focused on detecting direct signals from myelin or from iron, which is enriched in the cell body of oligodendrocytes. Magnetisation Transfer-based techniques, for example, quantify the fraction of macromoleculebound water protons, and have been shown to relate strongly to myelination in a number of validation studies (Deloire-Grassin et al., 2000; Dousset et al., 1995, 1992). R2* mapping, on the other hand, quantifies local field distortion caused by iron, and has been confirmed as an iron marker by several validation studies (Langkammer et al., 2010; Sun et al., 2015). R1 has gained attention recently as a quantitative metric for myelination, and although its effectiveness as a WM myelin marker has not been directly tested, it has been shown to detect spatial distributions of myelin in grey matter (Lutti et al., 2014; Stüber et al., 2014). In addition to the development of new MR techniques, new statistical tools, such as joint inference permutation testing (Winkler et al., 2014, 2016), facilitate the integration of Magnetic Resonance Imaging (MRI) techniques to clarify the biological interpretation of MRI-measured effects in white matter.

**Figure 1:**
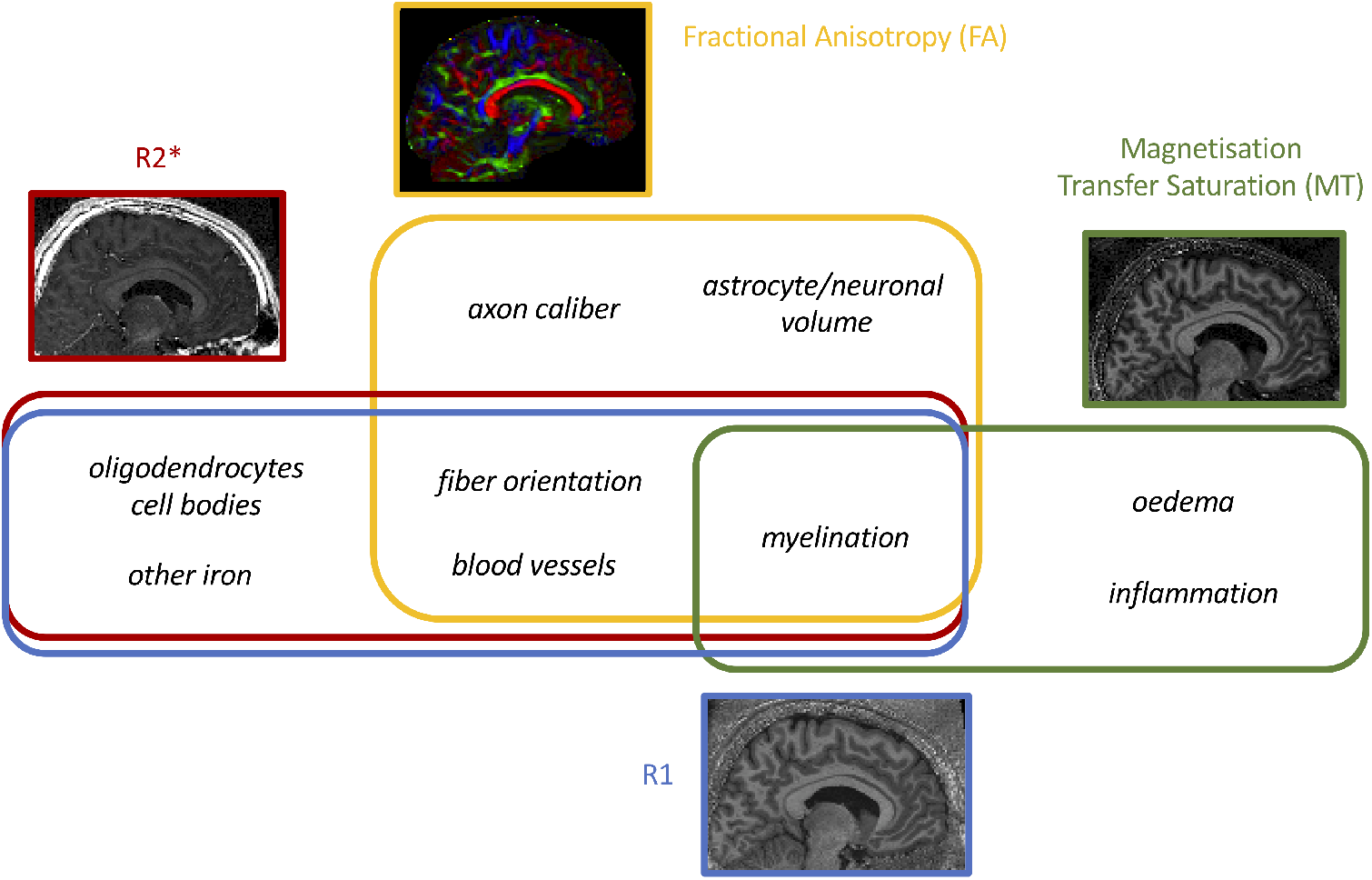
Each neuroimaging modality is sensitive, but not specific, to different features of the biological tissue. This study aimed to use multiple MR modalities that are sensitive to myelin, but measure different biophysical properties of white matter.

Applying these approaches to studying WM microstructural techniques could be helpful for clarifying the mechanisms behind WM-behaviour relationships. In particular, using MRI modalities that are sensitive to different biophysical tissue properties could disentangle whether myelination, oligodendrocytes, or fiber orientation, or a combination of them, are key in driving reported FA-behaviour correlations. In turn, if all WM-behaviour relationships are driven by a common biological mechanism, then establishing recurrent multimodal patterns that correlate with behaviour could uncover it, with powerful implications for future studies looking at WM-behaviour relationships and biomarker development.

To tackle these open questions regarding WM-behaviour relationships, we set out to:

1. Perform confirmatory, pre-registered testing of FA-behaviour relationships.
2. Perform pre-registered testing of relationships between behaviour and microstructural imaging across neuroimaging modalities.
3. Identify multimodal microstructural signatures which may provide insights into the underlying biology of WM-behaviour relationships.

### Participants

Figure 2 summarises the study design. 50 healthy participants (25 female; aged 18-38 years, mean 26.2 years, median 26 years) underwent a single session of behavioural testing and MRI on the same day. As there is limited literature on the sample sizes needed to robustly detect cross-sectional correlations, our target sample size was based on previous work which had informed our hypotheses (n=20 for DSST (Metzler-Baddeley et al., 2012), n=21 for AFT, as the average sample size in the studies reviewed by (Gooijers and Swinnen, 2014), and n=26 for TOJ (Husain et al., 2011)). Studies reporting positive results may underestimate the necessary sample sizes (Button et al., 2013), so we doubled the sample size reported from the literature, thus bringing our sample size in line with a report recommending samples sizes between n=20 and n=40 for studies on FA (De Santis et al., 2014).

**Figure 2:**
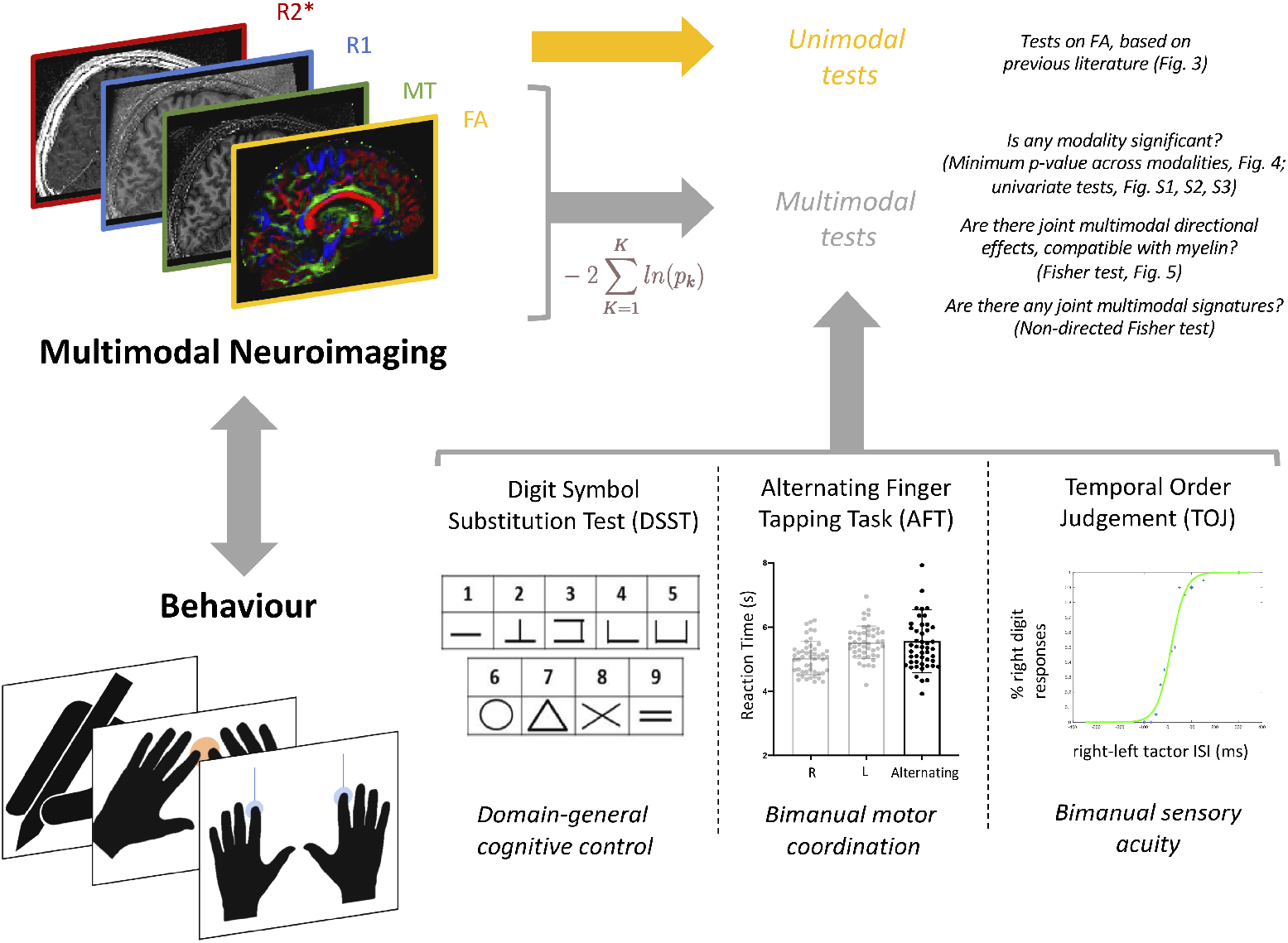
Study design and summary of MRI and behavioural data acquired.

All participants were self-assessed right-handed and their handedness was further assessed through the Edinburgh Handedness Inventory (Oldfield, 1971) (score range 60-100, mean 87.2, median 90). All participants were screened for MRI safety, received monetary compensation for their participation, and gave their informed consent to participate in this study. All study procedures followed the Declaration of Helsinki, and were reviewed and approved by the local ethics committee at the University of Oxford.

### Preregistration

Details of the task data collection and analysis plans were preregistered on the Open Science Framework website (full pre-registration available here: https://osf.io/ar7zs/). In brief, the pre-registration covered hypotheses and aims of the project, including which behavioural measures, MR metrics and regions of interest to use, while analytical details were decided separately after data collection.

We report here relevant text from the pre-registration: “Overall aim: testing whether previously reported correlations between behavioural measures and fractional anisotropy (FA) measures in long-range projections obtained using diffusion-weighted magnetic resonance imaging (dw-MRI) are related to indices of myelin content obtained using novel quantitative magnetic resonance imaging (qMRI) protocols. To this end, we aim to replicate a sample of previous studies, and extend these FA/behaviour analyses to myelin qMRI/behaviour analyses”.

Specific brain/behaviour predictions were made for each task, listed in the analysis section below.

### Behavioural tasks

A set of behavioural tasks was selected to build on prior studies reporting relationships between behaviour and WM microstructure.

The presence of FA-behaviour relationships has been particularly clear for the corpus callosum and for the cingulum. The cingulum has been often implicated in cognitive control (Bathelt et al., 2019), and cingulum FA has been found to strongly correlate with performance on neuropsychological tasks (Metzler-Baddeley et al., 2012). The corpus callosum, on the other hand, allows the nodes of the motor network in each hemisphere to communicate with one another, and both positive and negative relationships have been widely reported between callosal FA and various types of bimanual performance ((Johansen-Berg et al., 2007; Muetzel et al., 2008; Sullivan et al., 2001) and (Gooijers and Swinnen, 2014) for a comprehensive review of callosal-bimanual behaviour relationships).

FA-behaviour relationships have also been thoroughly explored in behavioural paradigms beyond the motor system. As mentioned above, bimanual motor performance has been the subject of much literature, and so has bilateral sensory processing. In the visual domain, topographic organisation and visuospatial capacity have both been shown to relate to callosal microstructure (Saenz and Fine, 2010; Todorow et al., 2014). In the auditory domain, relationships have been established between perceptual acuity and WM microstructure, although mostly in pathology (Husain et al., 2011; Lin et al., 2008; Wang et al., 2019). While there have been no previous studies on WM relationships with somatosensory acuity, it would be logical to expect a similar relationship between somatosensory perceptual acuity and microstructure of WM in relevant tracts.

Specifically, we assessed three task domains:

1. testing for a relationship between callosal FA and bimanual motor performance using the Alternating Finger Tapping task aimed to directly replicate a series of previous studies (reviewed by (Gooijers and Swinnen, 2014));
2. testing for a relationship between cingulum FA and performance using the Digit Symbol Substitution Test (Metzler-Baddeley et al., 2012). Previous findings for this task were only reported in older adults (age range: 53 to 93, mean age: 74 (Metzler-Baddeley et al., 2012)), accounting for confounding effects of age. Here, to maintain comparability to the other tasks studied, we tested a younger population.
3. testing for a relationship between FA in somatosensory tracts and somatosensory perceptual acuity using the Temporal Order Judgement Task aimed to extend previous findings in the visual and auditory domain, to the sensory system.

These three tasks are described in detail below.

### Digit Symbol Substitution Test (DSST)

A paper-based Digit Symbol Substitution Test (DSST) was conducted as per https://healthabc.nia.nih. govsitesdefaultfilesdsst 0.pdf. After training on substituting 10 digits for symbols, participants were asked to sequentially fill in the remaining 90 symboldigit boxes in 90 seconds.

### Analysis of the DSST

The score was calculated as the total number of symbols filled in correctly by the end of the task. Two participants were identified as outliers (>3 SD away from the mean) and thus excluded from further analyses.

### Alternating Finger Tapping (AFT) task

The finger tapping task aimed to test the participants’ bimanual coordination. The task was based on (Muetzel et al., 2008) and (Pelletier et al., 1993) and ran as follows: three blocks were repeated four times (the first one for training purposes): during the first block, participants were asked to tap their right index finger on a buttonbox (Current Designs, Inc., Philadelphia, PA) 30 times, as fast as they could (right monomanual condition); during the second block, participants were asked to tap their left index finger (left monomanual condition); during the third block, participants were asked to alternate between right and left index finger button presses (bimanual condition). For each block, after the 30 button presses were finished, the total elapsed time was fed back on the computer screen. The experimenter inspected the participant movement by eye to ensure they were correctly switching between fingers and that they were moving the finger rather than the hand. Participant posture and hand position was carefully kept constant throughout all blocks. One participant did not carry out the AFT due to a hardware problem.

### Analysis of the AFT task

Alternating Finger Condition (AFC) duration was extracted, i.e. average total time needed for 30 taps on the alternating finger condition (Muetzel et al., 2008). Two participants were identified as outliers (>3 SD away from the mean) and thus excluded from further analyses. Total time needed for 30 taps on the monomanual conditions was used as a covariate in group-level analyses, (Pelletier et al., 1993), together with age and gender.

### Temporal Order Judgement (TOJ) task

The Temporal Order Judgement (TOJ) task aimed to test participants’ capacity to discriminate between two closely timed tactile stimuli delivered to the fingertips. The task was based on a previous investigation of the functional activity associated with such behaviour (Kolasinski et al., 2016) and ran as follows. A PC running a PsychoPy script delivered, via a USB 6501 card (National Instruments) and an amplifier (Tactamp, Dancer Design), two asynchronous pulses to two vibrotactile stimulators (also known as tactors, Dancer Design) positioned within holes in a foam pad. The participant was asked to keep their hands relaxed on the foam pad, with their index fingers gently lying on the tactors. A piece of cardboard was used to block visual input from the tactors; similarly, headphones playing low levels of pink noise were used to block the auditory input from the tactors. Participants performed a two alternative forced choice (2AFC) task and were asked to press on one of two foot pedals, depending on the side of the pulse that they thought had come first. Participants were asked to respond within 2 seconds. If they did not respond within this time then no response was recorded and a new trial was started. They were also instructed that if it was hard to judge which pulse came first, they should just make their best guess. Intervals between pulses ranged from 0 to 300 ms. The task featured a practice session with 10 trials and a full session with 280 trials, for a total duration of roughly 12 minutes.

### Analaysis of the TOJ task

After trials with no response were discarded, the number of correct pedal responses were plotted as a function of interstimulation interval and a logistic regression was fitted to the data. At this stage, six participants were excluded as the logistic regression failed to fit the data correctly. The slope of the curve and the Just Noticeable Difference (JND) were used as key metrics of performance on the task (Kolasinski et al., 2016; Shore et al., 2005).

### MRI data collection

Magnetic Resonance Imaging (MRI) data were collected with a 3.0-T Prisma Magnetom Siemens scanner, software version VE11C (Siemens Medical Systems, Erlangen, Germany). Participants were asked to keep their head still and to wear earplugs during scanning in order to reduce the impact of MRI-related noise. The sequences were collected as follows: T1-weighted structural imaging (T1w), resting-state fMRI (rs-fMRI), Multi-Parameter Mapping (MPM) and Diffusion-Weighted Imaging (DWI). MRI scan pre-processing, analysis and statistical comparisons were performed using FMRIB Software Library (FSL, v6.0), except for the MPM quantitative map estimation step which was carried out using the hMRI toolbox implemented in Matlab-based SPM, as described in (Tabelow et al., 2019).

The T1w sequence had a TR of 1900 ms, TE of 3.96 ms, a 1mm isotropic resolution and a large Field of View (FOV, 256 mm^3^) to allow for the nose to be included in the image and thus facilitate neuronavigation later on in the paradigm. The sequence used GRAPPA with an acceleration factor of 2.

The diffusion-weighted Echo-planar imaging (EPI)sequence had TR=3070 ms, TE=85 ms, FOV=204mm^3^, voxel size=1.5mm isotropic, multiband factor of 4. Diffusion scans were collected for two b-values (500 and 2000 *s/mm^2^),* over 281 directions. An additional 23 volumes were acquired at b=0, 15 in AP phase-encoding direction and 8 in the PA phase-encoding direction.

The MPM protocol (as per (Weiskopf et al., 2013)) included three multiecho 3D FLASH (fast low-angle shot) scans with varying acquisition parameters, one RF transmit field map (B1+map) and one static magnetic (B0) field map scan, for a total acquisition time of roughly 22 minutes. To correct for interscan motion, position-specific receive coil sensitivity field maps, matched in FOV to the MPM scans, were calculated and corrected for (Papp et al., 2016). The three types of FLASH scans were designed to be predominantly T1-, PD-, or MT-weighted by changing the flip angle and the presence of a pre-pulse: 8 echoes were predominantly Proton Density-weighted (TR = 25ms; flip angle = 6 degrees; TE = 2.3-18.4ms), 8 echoes were predominantly T1-weighted (TR = 25ms; flip angle = 21 degrees; TE = 2.3-18.4ms) and 6 echoes were predominantly Magnetisation Transfer-weighted (MTw, TR = 25ms; flip angle = 6 degrees; TE = 2.3-13.8ms). For MTw scans, excitation was preceded by off-resonance Gaussian MT pulse of 4 ms duration, flip angle of 220 degrees, 2 kHz frequency offset from water resonance. All FLASH scans had 1 mm isotropic resolution and field of view (FOV) of 256×224×176 mm. The B1 map was acquired through an EPI-based sequence featuring spin and stimulated echoes (SE and STE) with 11 nominal flip angles, FOV of 192×192×256 mm and TR of 500 ms. The TE was 37.06 ms, and the mixing time was 33.8 ms. The B0 map was acquired to correct the B1+ map for distortions due to off-resonance effects. The B0 map sequence had a TR of 1020.0 ms, first TE of 10 ms, second TE of 12.46 ms, field of view (FOV) of 192×192×256 mm and read-out bandwidth of 260 Hz/pixel.

### MRI preprocessing

A custom pipeline based on existing FSL tools (Smith et al., 2004) was developed for our diffusion sequence. The topup tool was run on average images of AP b0 volumes and PA b0 volumes. The resulting susceptibility-induced off-resonance field was used as an input for the eddy tool (Andersson and Sotiropoulos, 2016), which was run with options optimised for multiband diffusion data to correct for eddy currents and subject movement. To generate Fractional Anisotropy (FA) maps, a diffusion tensor model was fit to each voxel through DTIFIT.

Magnetisation Transfer saturation (MT), R1 and R2* quantitative maps were estimated through the hMRI toolbox (Tabelow et al., 2019), with default settings including ESTATICS modelling (Weiskopf et al., 2014). In order to register MPM volumes to FA volumes, we used the following steps. Boundary-Based Registration was used to calculate a DWI-to-T1w registration using preprocessed b0 images (with high tissue boundary contrast). A customised pipeline was used to apply the fslreorient2std tool to the MPM maps and register them to T1w space. At this stage, 1 participant was excluded as the MPM-derived maps were heavily corrupted due to movement artefacts; 1 participant was excluded due to lower quality signal in the MPM scan, which resulted in poor registration with other modalities. Once registration matrices for MPM-T1w and DWI-T1w were calculated, they were inverted, concatenated and applied as needed to bring MPM volumes into DWI space with minimal interpolation. Registrations were assessed manually and one participant was excluded due to poor registration across all analyses.

### MRI analysis

To bring all volumes into a common space, native FA volumes were skeletonised with Tract-Based Spatial Statistics (TBSS (Smith et al., 2006)), and the skeletonisation transforms were subsequently applied to MPM-to-DWI registered volumes. Group-level analyses were then conducted in skeleton space for all data.

All behavioural performance measures were normalised (through z-scoring, or rank-based inverse-normal transformation if not normally distributed) and correlations between MRI metrics and behaviour were assessed for each behavioural measure separately.

Relevant text from the preregistered analysis plan is as follows:

Cingulum and DSST: “We aim to replicate a reported relationship between [...] number of substituted digits in the Digit Substitution test and cingulum FA (Metzler-Baddeley et al., 2012) [...], and to extend the protocol to investigate qMRI [...]/behaviour relationships.”

Callosum and AFT: “We aim to replicate a reported relationship between callosal FA and AFC duration in the finger tapping task (Sullivan et al. 2001; Muetzel et al., 2008). We further aim to test for a relationship between myelin metrics in the corpus callosum and AFC duration”

Sensorimotor tracts and TOJ: “Performance on the temporal order judgement task is not associated with integrity of a single specific white matter tract, but rather with a set of tracts involving multiple sensorimotor areas. Accordingly, we plan to run exploratory analyses across the whole brain, testing for associations between JND/slope values and FA/qMRI.”

Covariates of age, sex, and performance on control tasks (unimanual finger tapping speed for the AFT, and visuomotor speed for DSST) were included. For each behavioural assay, voxelwise analyses were restricted to voxels within a predefined anatomical mask chosen from standard atlases included in FSL and based on the a priori hypotheses: a cingulum mask for DSST, a callosal mask for AFT and a mask of cortico-cortical and ascending sensorimotor tracts for TOJ. The masks were derived from the JHU ICBM-DTI-81 Atlas, the JHU White-Matter Tractography Atlas and the Human Sensorimotor Tracts Atlas, respectively.

Within these masks, analyses were conducted with voxelwise maps of FA, MT, R1 and R2*. Voxelwise inference across these MRI modalities, testing for correlations between each MRI modality and behavioural measures, was performed using the Permutation Analysis of Linear Models (PALM) tool (Winkler et al., 2014). Cluster-wise inference was conducted to control familywise error over the image. A cluster-forming threshold of t>1.7 (equivalent to p<0.05, based on the degrees of freedom) was used in all instances, at the 5% familywise error level.

### Unimodal tests of FA

For unimodal hypotheses on FA, we reported the univariate results for correlations between FA and behaviour.

### Multimodal tests

For multimodal hypotheses, voxelwise inference using Non-Parametric Combination (NPC), as implemented in PALM (Winkler et al. 2016), was used to produce two types of inferences. (1) Correcting over modalities allowed us to ask whether *any individual modality* correlates with behaviour; (2) Combining over modalities allowed us to ask whether *any combination of modalities* correlates with behaviour.

For approach (1), we conducted cluster-wise inference on each modality separately, with familywise error controlled over the image and the *K* modalities. For each voxel, we reported the minimum image/modality-corrected cluster p-value across modalities.

For approach (2), combining evidence of effects over *K* modalities, we used Fisher’s p-value combining method at each voxel:

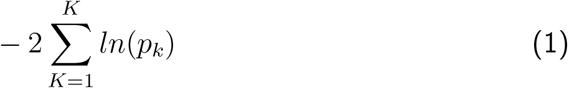

With this approach, evidence can be assessed for either directional or non-directional effects: combining one-sided p-values (based on prior expected directions of effects) will test for directional effects; combining two-sided p-valueswill provide sensitivity to non-directional effects (i.e., combination of either direction) as well. Here, a directional Fisher test, testing for positive effects across all modalities, was used to test for putative myelin signatures.

### Simulation-based post-hoc power calculations for combined multimodal tests

A comprehensive power analysis for cluster-wise inference that accounts for the spatially-varying dependence among imaging modalities is beyond the scope of this work. However, so as to provide a rough indication of power for future studies of multimodal microstructural imaging, we conducted univariate simulation-based power calculations for the combined multimodal (Fisher) tests. Pearson correlations for each modality-behaviour pair were recorded at the location of the peak voxel in the Fisher test inference map. In each simulation, a Gaussian random vector of behavioural and imaging values were generated with the specified correlation induced between the behaviour and each imaging value. We then tested whether the null hypothesis for each simulation would be rejected under a Fisher test with alpha set at 0.001. Power was then calculated as the percentage of tests rejecting the null hypothesis across all simulations. For each WM-behaviour correlation, power was calculated for samples sizes ranging from 10 to 300 subjects. While this approach may be optimistic because of using a peak voxel to measure effect sizes, it probably is conservative since it represents power at a single voxel and does not reflect the sensitivity gained through cluster inference.

## Results

We first used unimodal analyses to test for correlations between DWI-derived FA and behaviour, based on previously reported literature (Figure 3). No relationships were found between behaviour and FA within tracts of interest for either TOJ or DSST (TOJ: peak p_corr_=0.08; DSST: peak p_corr_=0.49). For AFT, a significant correlation was found between callosal FA and AFT performance (peak p_corr_=0.016).

**Figure 3:**
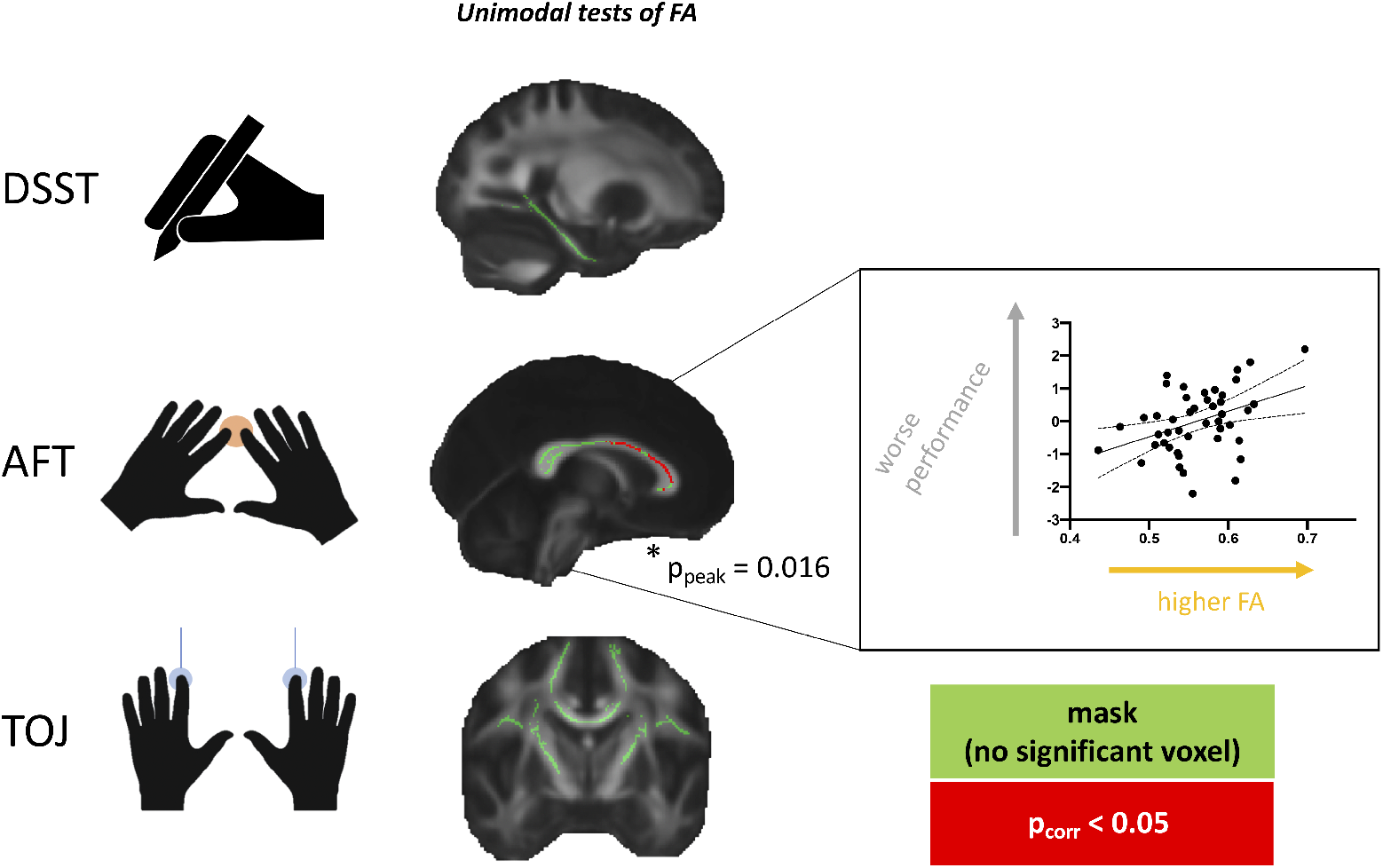
FA and behaviour. Unimodal relationships between FA and behaviour were tested across anatomical masks (shown in green) that were selected for each task. Results highlight that the Alternating Finger Tapping task (AFT), but not Temporal Order Judgement task (TOJ) and Digit Symbol Substitution Test (DSST) has a significant relationship with FA (red cluster shows voxels with corrected p-values below 0.05). Within that cluster, mean FA is extracted for each subject and plotted against performance in the scatterplot (with line of best fit and 95% confidence bands), that is for visual assessment of the correlation, rather than for statistical inference.

We then performed multimodal tests, testing whether *any individual* modality (FA, MT, R1 or R2*) strongly correlated with behaviour (Figure 4), by considering p-values across both voxels and modalities for each WM-behaviour relationship. No relationships were found between behaviour and multimodal MRI metrics within tracts of interest for either TOJ or AFT (TOJ: peak p_corr_=0.339; or AFT: peak p_corr_=0.09). For DSST, a significant correlation was found between parahippocampal cingulum and DSST (peak p_corr_=0.038), driven entirely by R1 (only modality with any voxel of p_corr_ <0.05, Figure 4).

**Figure 4:**
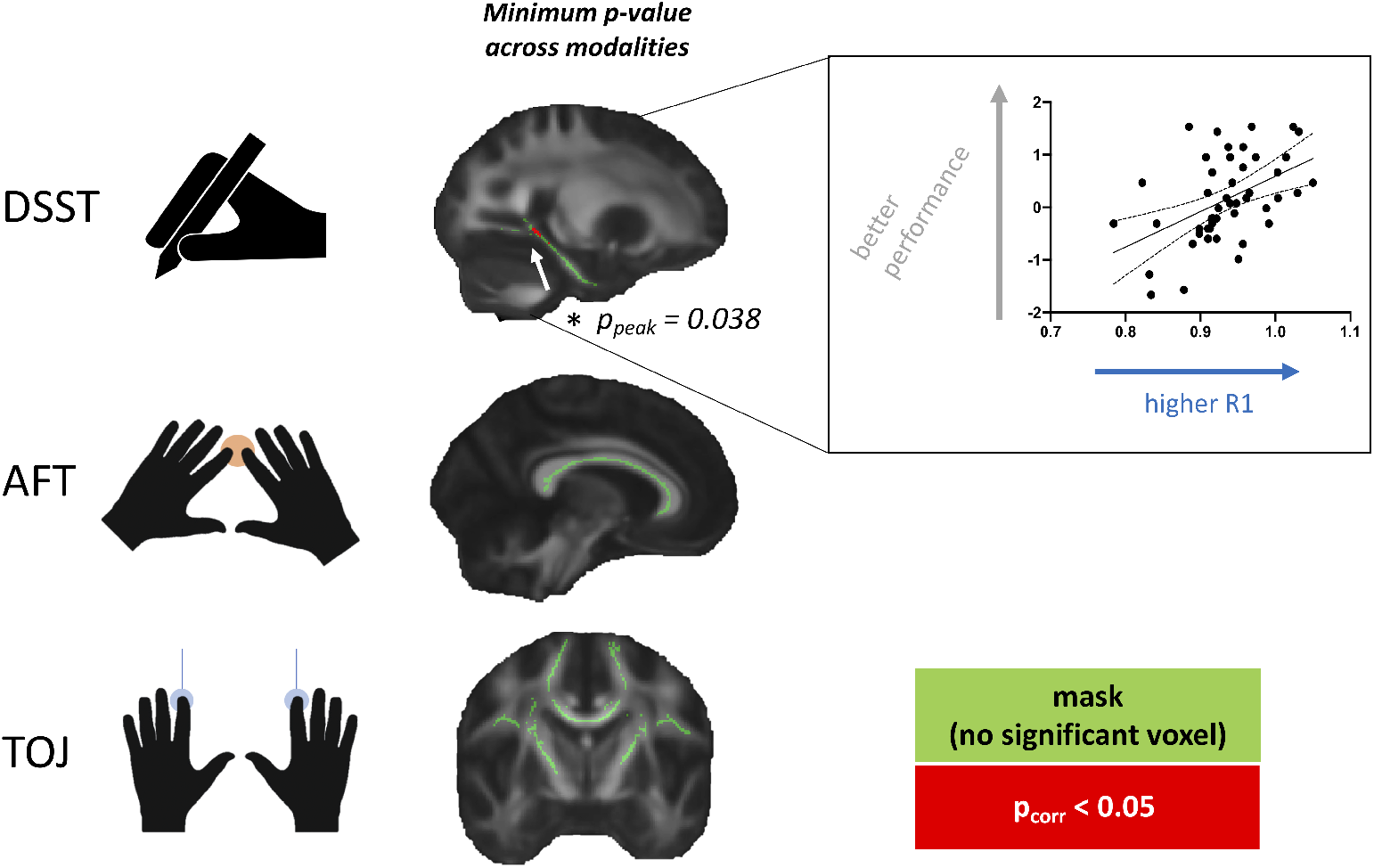
Multimodal microstructural imaging and behaviour. Multimodal relationships between behaviour and individual MRI metrics (FA, MT, R1 and R2*) across Digit Symbol Substitution Test (DSST), Alternating Finger Tapping task (AFT) and Temporal Order Judgement task (TOJ). Only the DSST has a significant relationship with cingulum WM, driven by R1, when considering FWER-corrected p-values (red cluster shows voxels with corrected p-values below 0.05). Within that cluster, mean R1 is extracted for each subject and plotted against performance in the scatterplot (with line of best fit and 95% confidence bands), that is for visual assessment of the correlation, rather than for statistical inference.

While single-modality tests allow to identify strong correlations with a particular modality, they cannot identify combined trends across modalities, which can be particularly informative of the underlying biology. For instance, a positive trend across all modalities considered here (which are known to positively correlate with myelin content of the tissue) would indicate that tissue myelination may be related to behavioural performance. Likewise, trends in discordant directions could also be informative, as they could unveil multimodal signatures related to other biological tissue properties such as vasculature and fiber orientation.

Fisher tests were used to detect combined multimodal trends between behavioural measures and MRI metrics (FA, MT, R1 and R2*). With the usual (directed, positive) Fisher test (Figure 5, 2^nd^ column), no relationships were found between behaviour and multimodal MRI metrics within tracts of interest (TOJ: peak p_corr_=0.532; AFT: peak p_corr_=0.184; DSST: peak p_corr_=0.2). With a non-directed Fisher test (results not shown), once again no relationships were found between behaviour and multimodal MRI metrics within tracts of interest. (TOJ: peak p_corr_=0.82; AFT: peak p_corr_=0.11; DSST: peak p_corr_=0.29) Taken together, these two tests argue against the presence of consistent multimodal microstructural signatures related to myelination or to other biological tissue properties.

**Figure 5.**
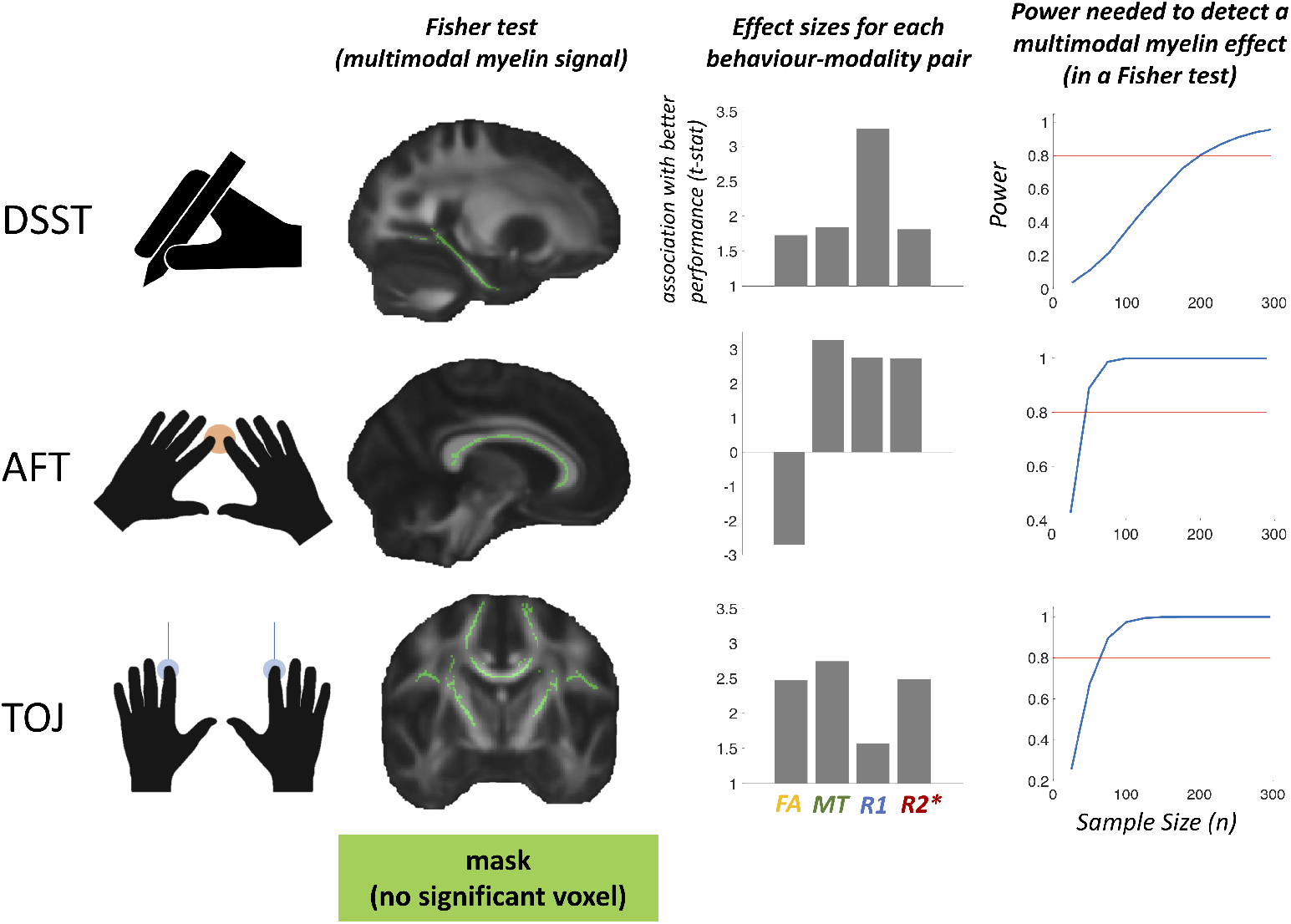
Lack of evidence for combined multimodal signatures. A Fisher test was used to search for multimodal microstructural signatures relating WM to behavior, but no significant effects were found (2^nd^ column). Effect sizes are reported for each modality-behaviour correlation, as measured by the top 5% t-statistic within peak Fisher clusters. This analysis was carried out to provide a clear visualisation of peak effect size for each pair of MR modality and behaviour, rather than for statistical inference (3^rd^ column). For each WM-behaviour correlation, we used a simulation-based approach to calculate sample sizes needed to reach 80% power (red line), given the observed effect sizes found in our pre-registered tests. Sample sizes needed to detect a combined multimodal effect vary from 190-200 participants for DSST, to 40-50 for AFT, to 60-70 for TOJ (4^th^ column).

The lack of a common microstructural signature is also apparent when considering the top 5^th^ percentile t-statistics (Figure 5, 3^rd^ column) and the t-statistics maps for each task (Figures S1, S2 and S3), where peaks are not consistent across modalities. This further confirms the negative Fisher tests, as there is no common trend across modalities within each group of WM-behaviour tests.

To aid future studies wishing to explore WM-behaviour correlations, and myelin-behaviour correlations in particular, we ran post-hoc simulation-based power analyses to identify the sample sizes needed to detect a combined multimodal effect through a Fisher test (Fig. 5, 4^th^ column). Based on the observed effect sizes, we find that sample sizes needed to detect a myelinbehaviour correlation across the 4 modalities in a directed Fisher test vary from 190-200 participants for DSST, to 40-50 for AFT, to 60-70 for TOJ.

For completeness, we also report analyses of this dataset using conventional univariate approaches, considering each modality separately (Figures S1, S2 and S3) and not correcting across modalities. We find that if each modalitybehaviour correlation was run as a separate analysis, each behaviour would show a correlation with at least one modality. Strikingly, different behaviours correlate most strongly with different modalities (DSST with R1 (Figure S1); AFT with FA and MT (Figure S2); TOJ with R2* (Figure S3)), thus strengthening the evidence against a common microstructural signature across behaviours.

## Discussion

Our first aim was to assess the robustness of relationships between white matter FA and behaviour across a range of behavioural tasks. We find a unimodal correlation between the structure of the corpus callosum FA and bimanual coordination, in accordance with previous literature (Bathelt et al., 2019; Johansen-Berg et al., 2007; Metzler-Baddeley et al., 2012; Muetzel et al., 2008; Sullivan et al., 2001). This confirms that individuals with lower callosal FA perform better in tasks requiring bimanual coordination. It also suggests that the extensive early literature on bimanual coordination and the corpus callosum (Gooijers and Swinnen, 2014) can be replicated, even with larger sample sizes and recent preprocessing pipelines.

However, a robust relationship between FA and behaviour was identified in only one out of three tasks considered here. This can be due to several reasons. One possible explanation is that effect sizes inferred from previous studies might be overinflated due to publication bias (Turner et al., 2008) and under-powered analyses (Button et al., 2013). However, it is worth noting that, of the three tasks considered here, only the FA-AFT experiment, which did successfully identify a FA-behaviour relationship, was a direct replication of a previous testing protocol. The other two tasks were designed as conceptual replications or extensions, but did not precisely replicate experimental conditions and analysis steps. For instance, our analyses employed Tract-Based Spatial Statistics (Smith et al., 2006), as well as recently developed preprocessing tools (Andersson and Sotiropoulos, 2016), both of which differed from some of the studies we based our hypotheses on (Metzler-Baddeley et al., 2012). While our aim was not to perfectly replicate analyses from previous papers, it is possible that differences in preprocessing may be driving discrepancies between our FA results and the results from previous studies. In summary, the relationships between FA and behaviour that have been established may be robust and replicable, but the experimental and analytic conditions under which they occur needs clarification.

A second aim of the present study was to probe whether multimodal MR can provide useful insights on WM-behaviour relationships. We find that this is the case for at least one of the WM-behaviour relationships we tested: R1 correlates with DSST performance, such that individuals with higher R1 perform better in the DSST task requiring cognitive control. Higher R1 could reflect greater myelin, oligodendrocytes, vasculature or other iron-rich tissue components. In this case, multimodal analysis allowed identification of a WM-behaviour relationship that would have not been detected by an analysis focused on FA in isolation. This confirms that there is value in multimodal imaging, as some modalities may be more sensitive to the presence of a relationship than others.

A third aim was to test whether there are common multimodal microstructural patterns in WM-behaviour relationships, which may provide insights into the underlying biology. We fail to find robust evidence for multimodal effects and cross-modality signatures. Rather, we find that effect sizes and directionality of effect in the relationship between each modality and each behaviour are highly heterogeneous. This means that MR modalities in each tract not only show heterogeneity in how they relate to the same behaviour, but there is also variation as a function of which tract-behaviour correlation is being considered.

A key insight from the study is therefore that the relationship between WM and behaviour is highly varied. Given that each modality has a specific pattern of sensitivity to the underlying biology (Figure 1), the results suggest that different aspects of WM biology may be driving different WM-behaviour correlations. There are two prominent sources of biological heterogeneity in white matter, which are likely relevant to the results in this study.

One driver of heterogeneity may be at the level of myelination. We selected metrics that were all sensitive to the amount of myelin in an imaging voxel (Figure 1), predicting that if myelination were responsible for WM-behaviour relationships, a common multimodal pattern across all relationships would be identified. Such patterns were not found, arguing against myelination as a common driver. However, such reasoning might be overly simple-minded. Histological studies have increasingly highlighted the heterogeneity of features in the myelinated axon, which can vary independently of each other (Almeida and Lyons, 2017). For instance, we know that Nodes of Ranvier, myelin sheath thickness, myelin sheath length, and number of myelin sheaths, can all independently affect an axon’s physiological properties, which one would expect, in turn, to shape behaviour (Kaller et al., 2017). Varying these features might have differing effects on the overall amount of myelin in a given voxel meaning that the imaging metrics used might not be equally sensitive to all relevant features of the myelinated axon.

A second important driver of heterogeneity is non-myelin features of WM. As exemplified in Figure 1, while all sequences we used are sensitive to myelin, some are also sensitive to fiber orientation and neuronal volume (FA), and some are sensitive to iron and vasculature (R1 and R2*). Therefore, one possible interpretation of the data is that the relationship between AFT performance and the corpus callosum is highly influenced by fiber orientation, whereas the relationship between the DSST performance and the cingulum is shaped by vasculature. Previous studies highlighted that both fiber orientation (Chang et al., 2017; Wedeen et al., 2005) and vasculature (Licht et al., 2011; Rhyu et al., 2010; Thomas et al., 2016) are important for brain function, and our data thus draw further attention to the fact that these factors may be influential in WM-behaviour relationships.

These two factors combined may explain why there is no single aspect of WM that drives behaviour. Rather, our findings confirm that heterogeneity at the cellular level is reflected in variation in the relationship between neuroimaging markers and behaviour. Importantly, this emphasizes that there is no single modality or single combination of modalities which is optimal to study WM-behaviour relationships. In this respect, our study poses practical limits to the possibility of developing a one-size-fits-all approach to the investigation of white matter-behaviour relationships, due to their inherent diversity.

While this heterogeneity means it is not straightforward to predict which MR modality is best suited for each type of WM investigation, it also suggests that multimodal studies of WM should tailor their MR sequence protocols and analyses pipelines to privilege markers and statistical approaches that can test and compare biologically-grounded models. For example, with an appropriate acquisition sequence and a joint multimodal statistical framework, one might be able to test whether a given WM-behaviour correlation is driven by myelination, vasculature (Thomas et al., 2016), or connectivity (Sui et al., 2014). Such approaches are most likely to generate further insights into WM-behaviour relationships in the future.

One key limitation of the study is that the results cannot disentangle to what extent differences between WM tracts contribute to the observed diversity of WM-behaviour relationships. One could argue, for example, that our results demonstrate that FA is more important for WM-behaviour relationships involving the corpus callosum, whereas R1 is more important for understanding the cingulum, while MT/R2* are more important in investigations of the corticospinal tract. Because each of the behaviours we selected relates to a different WM tract, it is impossible to disentangle whether different kinds of behaviours are most strongly driven by different microstructural patterns, or whether there is neuroanatomical heterogeneity in the importance of different microstructural features of each tract. Although both are likely to matter, further studies relating individual tracts to multiple behaviours are required.

Moreover, an additional limitation of the study lies in the extent to which it was pre-registered. While our pre-registration covered hypotheses and aims, including behavioural measures, MR metrics and regions of interest, it is now increasingly being acknowledged that many analytical choices in neuroimaging can have a large influence on the final results (Nichols et al., 2017; Pervaiz et al., 2020), and are thus crucial for confirmatory analyses. Therefore, we recommend future studies to include sample size and details of their preprocessing and statistical modelling in their pre-registrations when appropriate.

The results also hold useful lessons for statistical aspects of future multimodal studies of WM. WM-behaviour correlations often have small effect sizes, and in our results we find that these effects are sometimes not detected when multiple hypotheses are tested concurrently. Testing for effects across modalities increases the false discovery rate proportionally to the number of modalities tested, and thus needs to be adequately corrected for in order to reach appropriate interpretations (Winkler et al., 2016). However, while multiple comparison correction has long been the gold standard statistical advice, multimodal brain imaging studies often do not report whether, and if so, how, correction for multiple comparisons was carried out (Bezukladova et al., 2020; Winston et al., 2020). Surprisingly, even gold standard guidelines in the field like COBIDAS do not report best practices for statistical reporting in multimodal imaging (Nichols et al., 2017), and many packages that support multi-modality statistical testing do not allow joint statistical tests, thus leaving room for needless analytic flexibility. Our results suggest there is a need for increased transparency in reporting of multimodal statistics, which statistical guidelines on multimodal imaging might facilitate in the future. In this respect, our results also add weight to previous calls to pre-register the modalities to be used in a given analysis (Picciotto, 2018), and to report all tested modalities in publications.

This aspect of statistics in multimodal studies also needs to be taken into account when assessing the power of a given analysis. When modalities are analysed separately, multimodal studies require multiple statistical tests across modalities. Therefore, for the same effect size, a study analysing multiple modalities may need more subjects to achieve the same power, and it is important to take this into account in power analyses. We thus recommend using larger sample sizes for multimodal compared to unimodal studies. Alternatively, another solution is to use non-parametric multivariate tests (Winkler et al., 2014, 2016) and/or dimensionality reduction techniques (Groves et al., 2011; Sui et al., 2014), in scenarios where multimodal data are available but the data set size is only powered for unimodal tests. While there is little literature on multimodal power analyses for cross-sectional studies using microstructural imaging, our results indicate that sample sizes of 40 to 200 may be required to detect joint multimodal effects through non-parametric multivariate tests.

In conclusion, these results highlight a broad heterogeneity in white matter’s relationship with behaviour. They also underscore the added value of multimodal imaging approaches, as different neuroimaging modalities might be best suited to detect different WM-behavior relationships. However, this added value needs to be weighed carefully against the need for more power and/or dimensionality reduction approaches in multimodal studies. Finally, the results effectively limit the possibility of developing a one-size-fits-all approach to study white matter, and suggest that different aspects of WM biology may be driving different WM-behaviour correlations.

## Supplementary Results

**Supplementary Figure 1:**
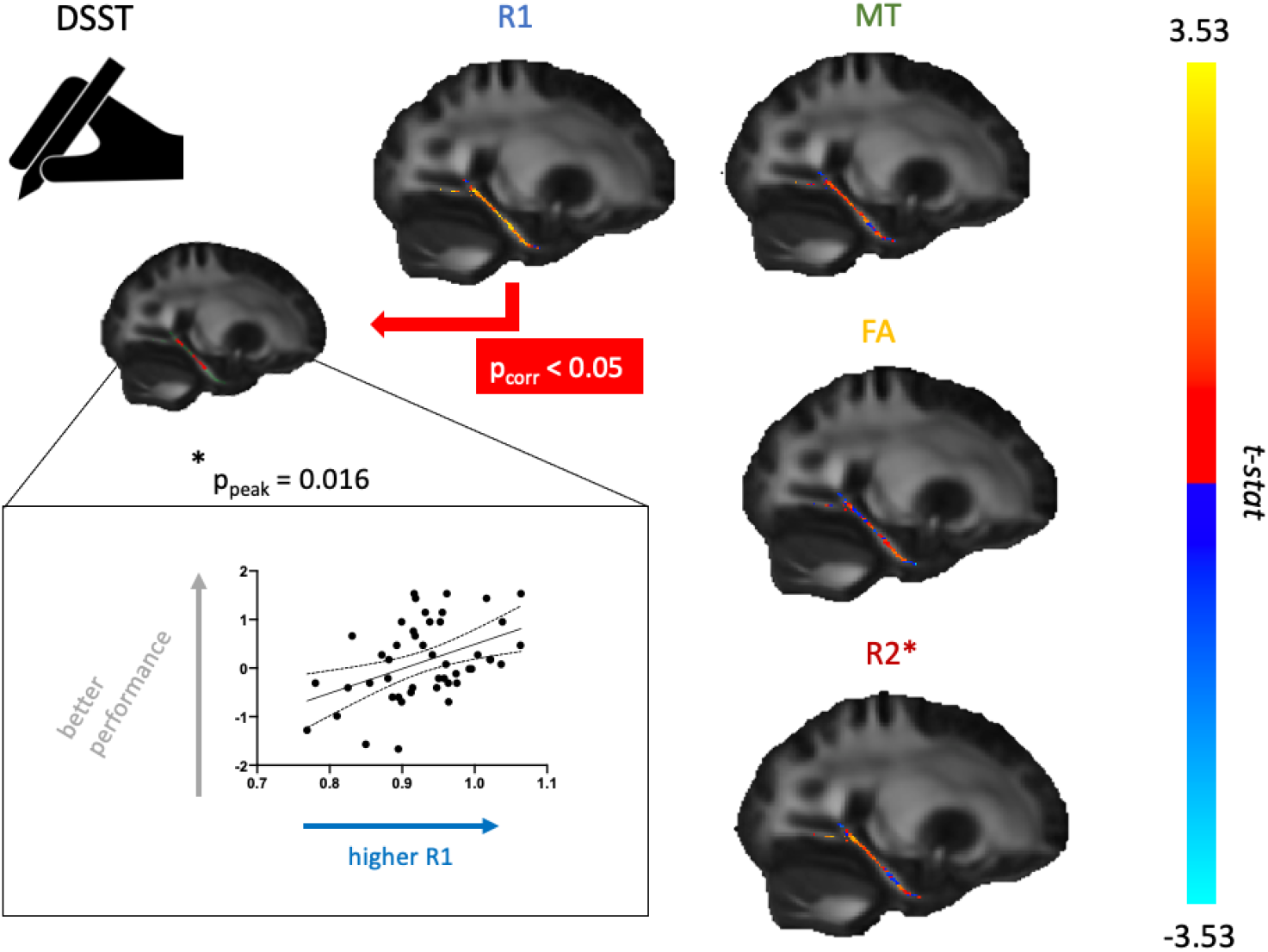
Correlation between DSST performance and cingulum microstructure, reported as univariate results. For each modality, unthresholded t-statistics are visualized according the colour bar (right). For R1 only, a cluster of voxels survived the threshold of p<0.05. Average R1 values within that cluster are shown against performance score in the scatterplot (with line of best fit and 95% confidence bands), which is presented for visualisation and is not used for statistical inference.

**Supplementary Figure 2:**
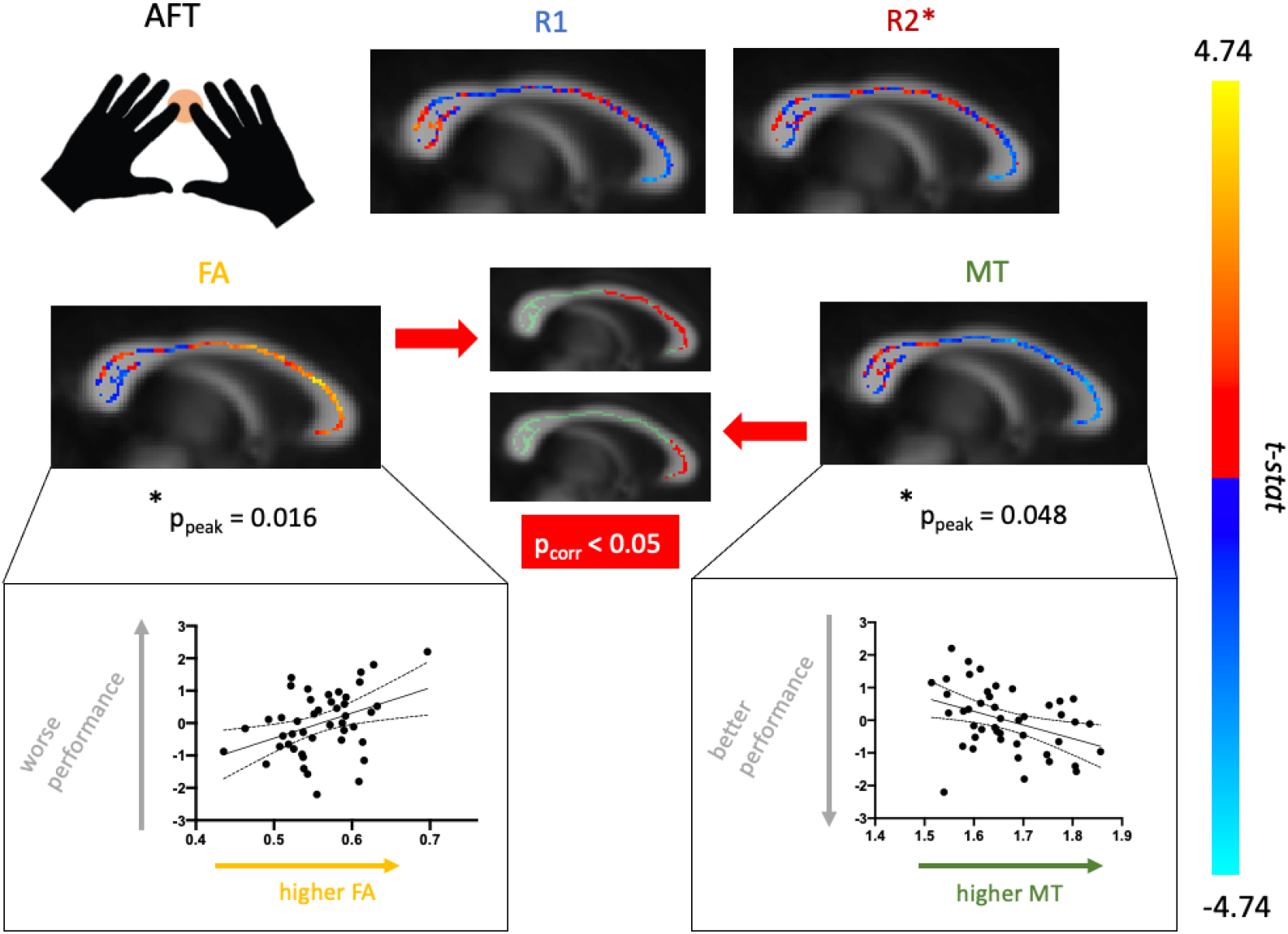
Correlation between AFT and callosal microstructure, reported as univariate results. For each modality, unthresholded t-statistics are visualized according the colour bar (right). For FA and MT, clusters of voxels survived the threshold of p<0.05. Average FA/MT values within that cluster are shown against performance score in the scatterplots (with line of best fit and 95% confidence bands), which are presented for visualisation and are not used for statistical inference.

**Supplementary Figure 3:**
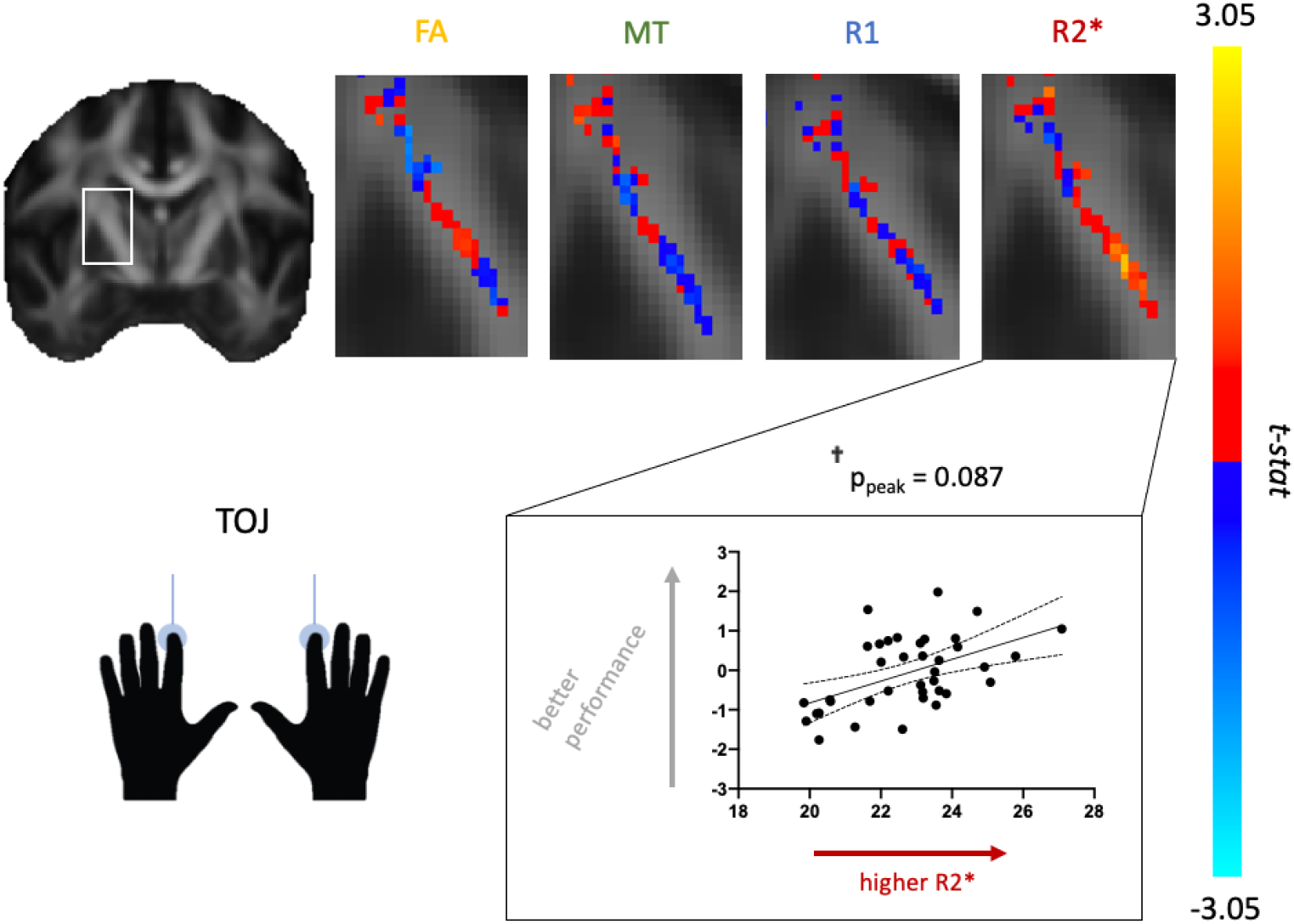
Correlation between TOJ performance and CST microstructure, reported as univariate results. For each modality, unthresholded t-statistics are visualized according to the colour bar (right). For R2* only, a cluster of voxels reached p=0.087. Average R2* values within that cluster are shown against performance score in the scatterplot (with line of best fit and 95% confidence bands), which is presented for visualisation and is not used for statistical inference.

## Acknowledgements

We are grateful to Ilona Lipp and Kate Watkins for their input on the manuscript. We thank Claudia Metzler-Baddeley for providing additional details on her previous experiments, which helped guide the DSST-Cingulum analyses. We would like to thank Juliet Semple, Nicola Aikin and Sebastian Rieger for their technical support and help with scanning participants; and Matthew Webster for wide-ranging help from IT to statistical issues. We acknowledge the IT-related support provided by David Flitney and Duncan Mortimer throughout the project.

## Data Availability Statement

Data used in this study is only available upon request due to data protection considerations.

## Funding

This work was supported by a PhD Studentship awarded to AL from the Wellcome Trust (109062/Z/15/Z) and by a Principal Research Fellowship from the Wellcome Trust to HJB (110027/Z/15/Z). CJS holds a Sir Henry Dale Fellowship, funded by the Wellcome Trust and the Royal Society (102584/Z/13/Z). OJW was funded by the Erasmus+ Programme by the European Commission. NE is a Wellcome Trust Doctoral student in Neuroscience at the University of Oxford [203730/Z/16/Z]. AJ was funded by the Dunhill Medical Trust (RPGF1810/93). The project was supported by the NIHR Oxford Health Biomedical Research Centre and the NIHR Oxford Biomedical Research Centre. The Wellcome Centre for Human Neuroimaging and the Wellcome Centre for Integrative Neuroimaging are supported by core funding from the Wellcome Trust [203147/Z/16/Z and 203139/Z/16/Z].

## Notes

### Competing Interest Statement

The authors have declared no competing interest.

